# Metabolic rates of prokaryotic microbes may inevitably rise with global warming

**DOI:** 10.1101/524264

**Authors:** Thomas P. Smith, Thomas J. H. Thomas, Bernardo García-Carreras, Sofía Sal, Gabriel Yvon-Durocher, Thomas Bell, Samrāt Pawar

## Abstract

Understanding how the metabolic rates of prokaryotes respond to temperature is fundamental to our understanding of how ecosystem functioning will be altered by climate change, as these micro-organisms are major contributors to global carbon efflux. Ecological metabolic theory suggests that species living at higher temperatures evolve higher growth rates than those in cooler niches due to thermodynamic constraints. Here, using a global prokaryotic dataset, we find that maximal growth rate at thermal optimum increases with temperature for mesophiles (temperature optima ≲ 45°C), but not thermophiles (≳ 45°C). Furthermore, short-term (within-day) thermal responses of prokaryotic metabolic rates are typically more sensitive to warming than those of eukaryotes. Given that climatic warming will mostly impact ecosystems in the mesophilic temperature range, we conclude that as microbial communities adapt to higher temperatures, their metabolic rates and therefore, carbon efflux, will inevitably rise. Using a mathematical model, we illustrate the potential global impacts of these findings.

## Introduction

A general understanding of how individual organisms respond to changing environmental temperature is necessary for predicting how populations, communities and ecosystems will respond to a changing climate ^1,2,3,4^. Because fundamental physiological rates of ectotherms are directly affected by environmental temperature ^3,5,6^, climatic warming may be expected to lead to ectotherm communities with higher metabolic rates on average ^3,7^. How environmental temperature drives metabolic rates of prokaryotes (bacteria and archaea) is of particular importance because they are globally ubiquitous, estimated to comprise up to half of the planet’s global biomass ^8^, and consume (respire) the majority of net primary production ^9,10^. Therefore, climate-driven changes in prokaryotic metabolic rates are expected to significantly alter ecosystem productivity, nutrient cycling, and carbon flux ^9,10,11,12,13,14^. Indeed, increased carbon efflux has been observed in experimental measures of soil CO_2_ loss to warming ^15,16^, as well as the responses of other microbial metabolic processes to increased temperature such as methanogenesis ^17^. However, whether the short-term (timescales of minutes to days) thermal responses of prokaryotes can be compensated by acclimation (physiological phenotypic plasticity) or longer-term (timescales of years or months, years or longer) evolutionary adaptation ^18,19,20^ is currently unclear. The most recent study to investigate this idea concluded that both short- and long-term responses of ecosystem-level heterotrophic respiration were similar^21^. However, this study quantified short-term responses by aggregating day-level carbon fluxes across sites, and did not have data on the direct respiratory contributions of prokaryotyes *per se*.

The short term, or “instantaneous” response of metabolic traits of individual organisms to changing temperature (the intra-specific thermal response) is typically unimodal, with the thermal performance curve (TPC) of the trait increasing with temperature up to a peak value (*T*_pk_), before decreasing as high temperature becomes detrimental to metabolic or cellular processes ^2,22^ (Fig. 1C). The *T*_pk_ for maximal population growth rate (a direct measure of fitness, often called the thermal optimum) is expected to correspond to the typical thermal environment in which the organism’s population has evolved (the long-term response) ^22,23^. The Hotter-is-Better (HiB) hypothesis posits that trait performance at *T*_pk_ (henceforth denoted by *P*_pk_) is also expected to increase inevitably in a similar manner to the short-term intra-specific response, because of the positive temperature-dependence of rate-limiting enzymes operating at their thermal optimum (a thermodynamic constraint), i.e. *P*_pk_ increases with *T*_pk_ (Fig 1A) ^22,23,24^. Thus this hypothesis essentially links the short term TPC of trait performance to the longer-term performance mediated by evolution. The HiB hypothesis is also implicit in the universal temperature dependence concept of the Metabolic Theory of Ecology (MTE) ^5,6,25^. However, whether the HiB hypothesis holds across organisms and environments is a question that is still debated ^24,26,27^. Deviations from a HiB pattern would indicate that thermodynamic constraints are compensated for by other mechanisms. In particular, an alternative hypothesis is that natural selection acts to override thermodynamic constraints, allowing peak trait performance and fitness to be, on average, equalised across different adaptation temperatures (Fig 1B) ^24^. Intermediate scenarios are also possible, where adaptation of optimal trait performance or fitness is only partially constrained thermodynamically (Fig 1C). Moreover, all enzyme families have a hard upper bound on the temperature at which they retain their functional integrity (which evolutionary changes cannot overcome), so very hot temperatures may in fact cause depressed maximal fitness (*P*_pk_ decreasing with *T*_pk_). Indeed, the existence of thermal constraints leading to an upper limit of prokaryotic growth rates has been shown recently ^28,29^. However a comparison of short- and long-term (HiB) responses of prokaryotic populations has never been made, nor the potential effects of responses at different time-scales on ecosystem fluxes studied.

**Figure 1:**
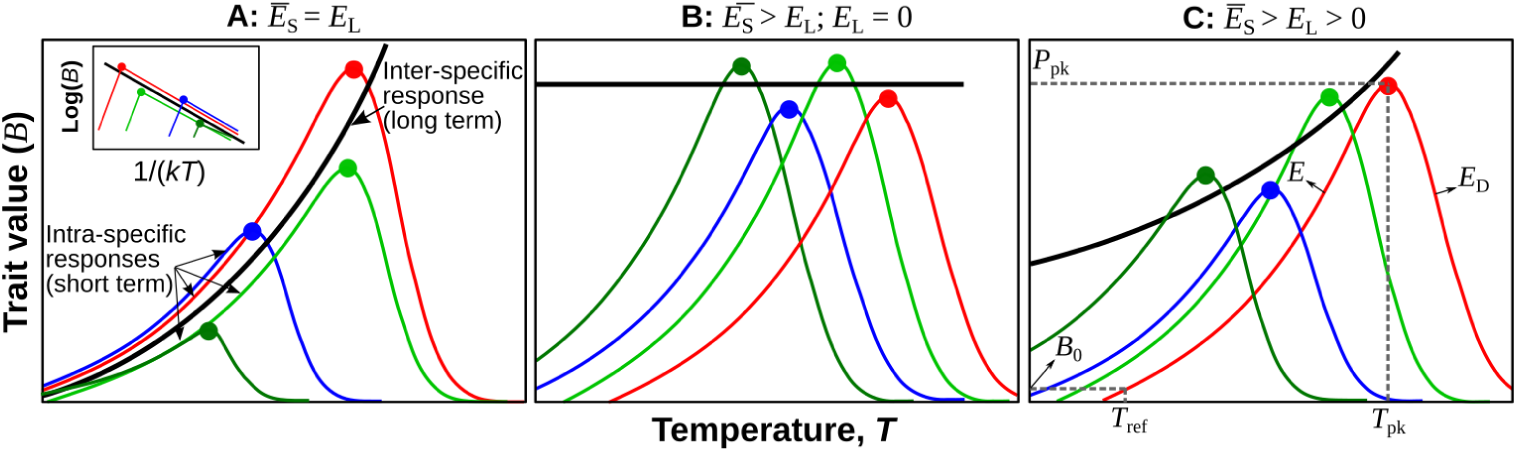
Three alternative hypotheses for the short- vs. long-term responses of thermal performance curves of a fitness-related metabolic trait in response to environmental warming. **A.** Hotter-is-Better: organisms adapt around a global, inter-specific, thermal constraint (black line, Boltzmann-Arrhenius fitted to intra-specific curve peaks), such that the average intra-specific (short-term) activation energy (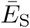) is statistically indistinguishable from the inter-specific (long-term) activation energy of the group of organisms (*E*_L_), and both are greater than zero. See methods for more details on the the definition and estimation of 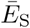 and *E*_L_, and the statistical methods used to differentiate between them. Note that each intra-specific TPC represents the short-term thermal response of each organism’s population. Inset panel illustrates how this would look in an Arrhenius plot. **B.** Equalisation of fitness: selection overrides thermodynamic constraints, such that trait performance at *T*_pk_ is on average the same (*E*_L_ = 0). Alternatively the same effect of *E*_L_ = 0 may occur due to or thermodynamic constraints on enzymes in fact restricting metabolic rate (and therefore fitness) at higher temperatures. **C.** Weak biochemical adaptation: an intermediate scenario where *E*_L_ *>* 0 but significantly less than 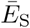. Panel C also illustrates the the Sharpe-Schoolfield TPC model parameters (eqn. 1, Methods).

Here we build and analyse a global dataset of TPCs in bacteria and archaea to quantify general patterns in both the short-term (intra-specific) response, and to test whether the HiB hypothesis holds (long-term, inter-specific response) within and across taxonomic and functional groups adapted to different temperatures (Fig. 1). These data go far beyond the scope of previous tests of the HiB hypothesis with or without microbes ^24^, covering practically the entire range of habitable global temperature niches (from bacteria isolated from Antarctic saline lakes at temperatures below 0°C, to a strain of methanogenic archaea able to proliferate at 122°C under high pressure) and the majority of the phylogenetic diversity of prokaryotes (spanning 9 bacterial phyla and the two major archaeal phyla, Euryarchaeota and Crenarchaeota; Fig. 3). In total we compare 542 growth rate TPCs and an additional 54 metabolic flux TPCs, spanning 482 unique prokaryotic strains.

## Results

### Adaptation To Culture Conditions

First, we compared each strain’s thermal optimum (*T*_pk_) with the temperature at which it was cultured (*T*_lab_) to determine whether the TPCs reflect adaptation to growth temperature. For both bacteria and archaea, we find a strong and significant (p *<* 0.00001) association between *T*_pk_ and *T*_lab_ (Fig. S1; bacteria *R*^2^ = 0.91, archaea *R*^2^ = 0.96) indicating that these strains are generally well-adapted to their culturing temperature. In both archaea and bacteria data subsets the *T*_pk_ vs *T*_lab_ line deviates significantly from a slope of 1 (bacteria slope = 0.87 *±* 0.04, archaea slope = 0.93 *±* 0.05) because *T*_pk_ tends to fall below culturing temperature at high temperatures (Fig. S1), suggesting a limit to thermal adaptation.

### Comparison of Short- and Long-term Thermal Responses

Next, we tested the HiB hypothesis by comparing the short-term (intra-specific) and long-term (inter-specific) thermal responses (see Fig. 1; Methods). If there is a universal thermodynamic constraint, peak fitness (*P*_pk_; *r*_max_ at *T*_pk_) across strains’ TPCs should increase with their respective *T*_pk_s (parameter *E*_L_; Fig. 1) at the same rate as *r*_max_ would increase with temperature (parameter *E*_S_), on average, within single strain’s TPC. Our analysis relies on *P*_pk_-*T*_pk_ pairs across strains because data within strains are largely lacking, and the HiB pattern is expected to apply across strains within monophyletic taxonomic groups (such as archaea and bacteria) ^24,30^. Analysing this relationship across 416 bacterial and 82 archaeal strains, we find that hotter is indeed better (HiB holds) across mesophiles (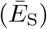 and *E*_L_ are *>* 0, and their 95% CIs overlap; Fig 2 and Table 1). However, this result does not extend to thermophiles, where instead fitness is on average invariant with respect to temperature. Thermophiles have evolved specific adaptations to extreme temperature stress, such as mechanisms to cope with increased membrane permeability at high temperatures ^31^ and thus adaptation to such niches may incur a fitness cost to thermophiles as seen in our results. This result is in concurrence with an investigation of the maximum growth rates of life on Earth, which found increases in microbial growth up to a peak before an attenuation of growth rates in warmer adapted organisms ^28,29^.

**Table 1:** Estimated mean short-term and long-term activation energies across archaea and bacteria, and test of the HiB hypothesis. Estimated mean *E*_S_ and *E*_L_ values (95% CI ranges in parentheses) for bacteria and archaea split by thermal niche (also see Fig 2). The last column indicates whether or not the HiB hypothesis is supported.

| Kingdom | Thermal Niche | $E_L$ | $\bar{E}_S$ | $E_L > 0$ | $\bar{E}_S \approx E_L$ | HiB |
| --- | --- | --- | --- | --- | --- | --- |
| Bacteria | Mesophile (n = 264) | 0.98 (0.75 – 1.25) | 0.87 (0.82 – 0.93) | TRUE | TRUE | TRUE |
|  | Thermophile (n = 114) | -0.07 (-0.26 – 0.12) | 0.85 (0.78 – 0.92) | FALSE | FALSE | FALSE |
| Archaea | Mesophile (n = 21) | 0.97 (0.69 – 2.26) | 0.60 (0.50 – 0.70) | TRUE | TRUE | TRUE |
|  | Thermophile (n = 60) | -0.09 (-0.21 – 0.17) | 1.11 (0.95 – 1.28) | FALSE | FALSE | FALSE |

**Figure 2:**
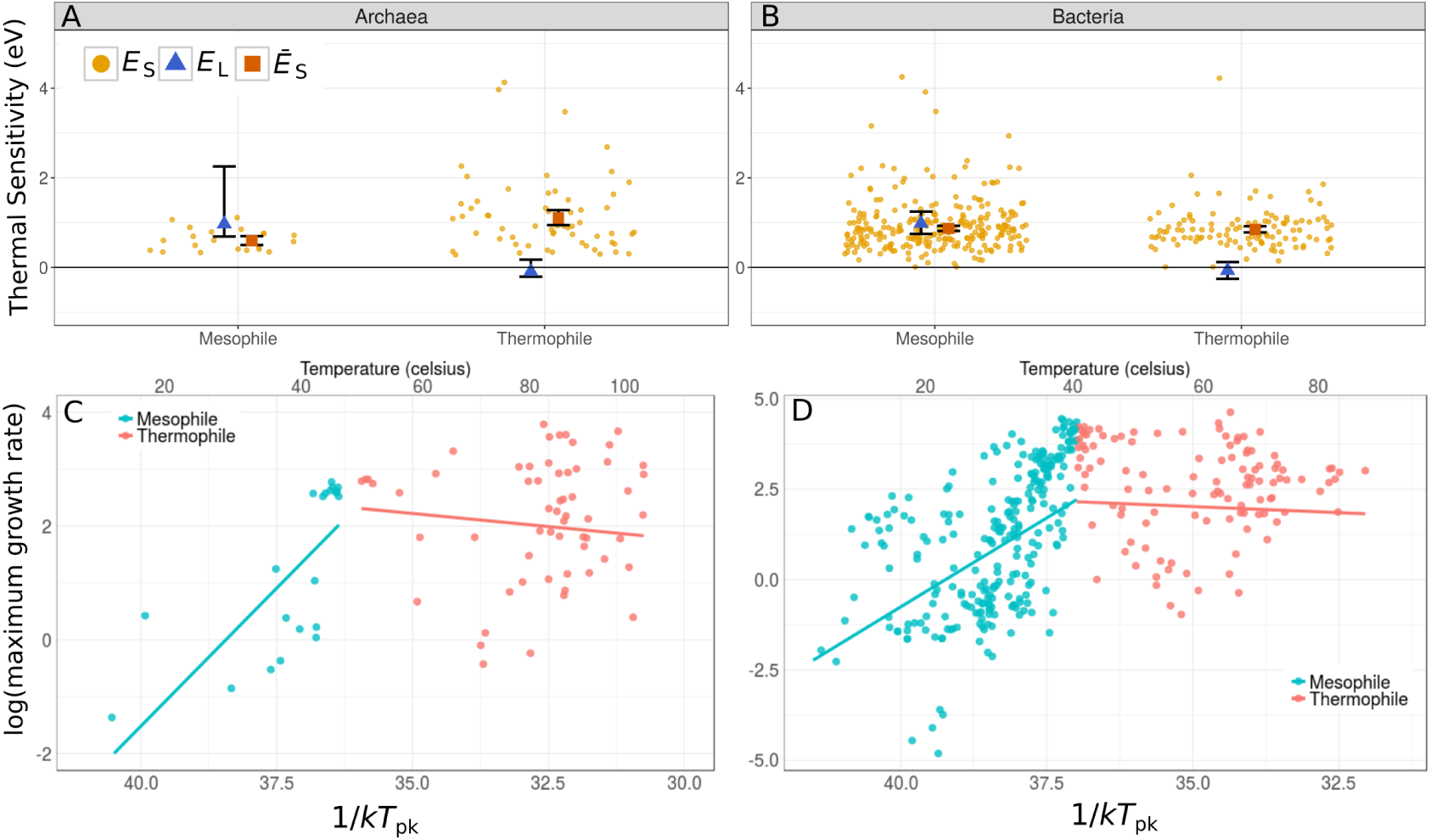
Patterns of short- and long-term thermal responses of growth rate (fitness) for archaea and bacteria. **A** and **B**: Activation energies (with 95% Confidence Intervals) from Boltzmann-Arrhenius model fits (*E*_L_, blue triangle) compared to mean activation energy (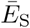, red square) of the intra-specific (short-term) thermal responses. Distribution of all the *E*_S_s is also shown (orange points). **C** and **D**: Arrhenius plots (x-axes inverted to aid visualisation) fitted to mesophile and thermophile sub-groups separately within bacteria and archaea, respectively. That is, the lines (the long-term thermal responses) are the Boltzmann-Arrhenius model fitted using weighted regression to mesophile and thermophile data separately, after determining the breakpoint (Methods). The HiB hypothesis is best-supported for the mesophile sub-group in both panels, while equalisation of fitness is best supported in the the thermophile sub-groups (Table 1).

**Figure 3:**
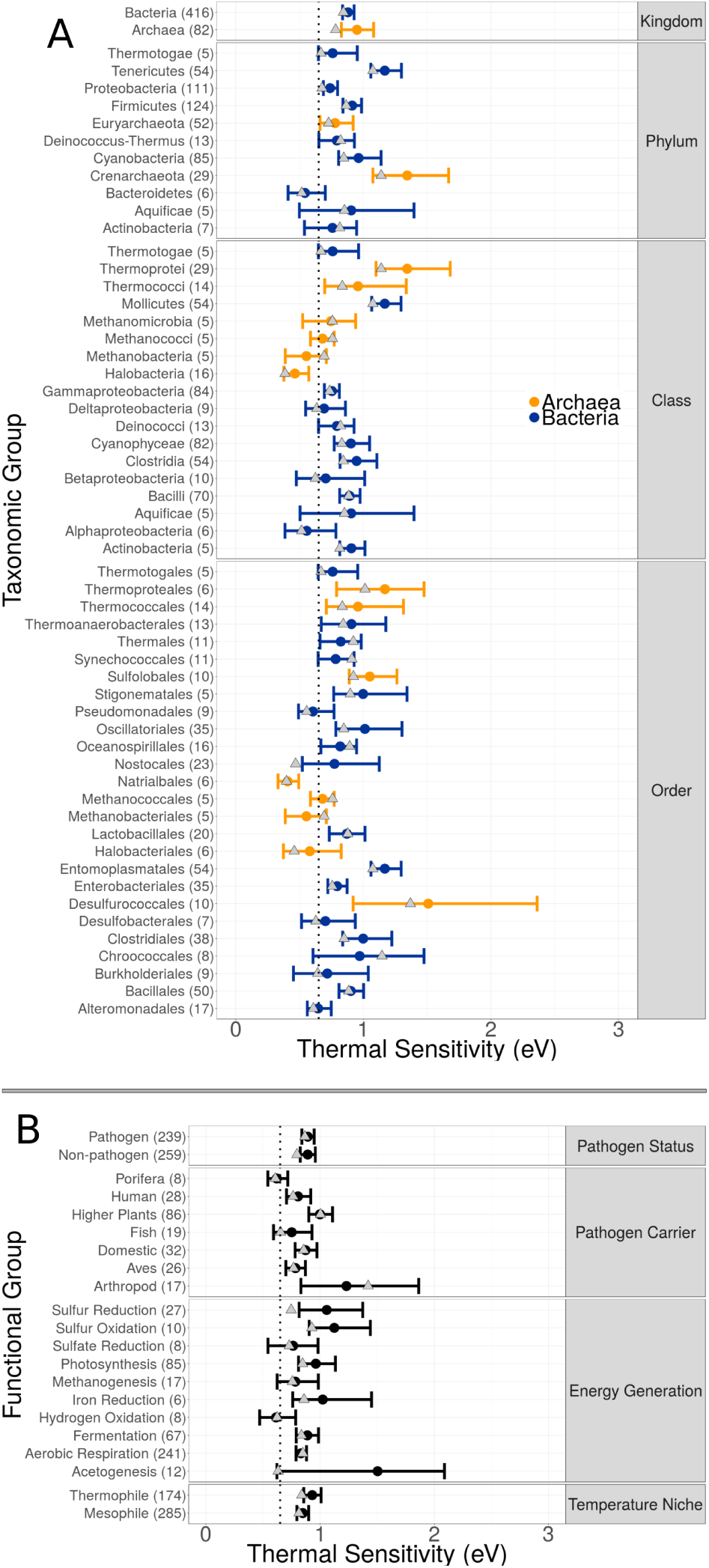
Variation in thermal sensitivity among prokaryotic groups. Comparison of intra-specific population growth rate activation energies (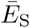) across taxonomic levels **(A)** and functional trait groupings **(B)**. Points and error bars represent weighted mean and 95% CIs of *E*_S_ for each group. Groups shown are those with at least five data points, the number in brackets indicates the number of data points from which 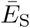 was calculated for each grouping. The dotted line marks 0.65eV, the mean *E* previously reported within the MTE framework. Grey triangles mark the median *E*_S_ for each group.

### Variation in Thermal Sensitivity

Under the MTE, the global (inter-specific) thermodynamic constraint is expected to center around 0.65eV ^5,6^. Mean intra-specific thermal sensitivities have been found to be very similar to this value, although the distribution is right-skewed with a median value of *∼*0.55eV ^2^. In contrast, we find that mean thermal sensitivities for both bacteria and archaea fall significantly above 0.65eV (Fig. 4, bacteria 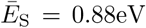; archaea 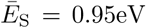). We observe the same right-skew in activation energies for prokaryotes as seen across other organisms and traits ^2^. Even after accounting for this skew by taking the median instead of the mean, activation energy still falls significantly above 0.65eV (bacteria median = 0.84eV, archaea median = 0.80eV; Fig. S2). Furthermore, we see a consistent pattern of median thermal sensitivity *>*0.65eV throughout the lower taxonomic groupings (Fig. 3A).

**Figure 4:**
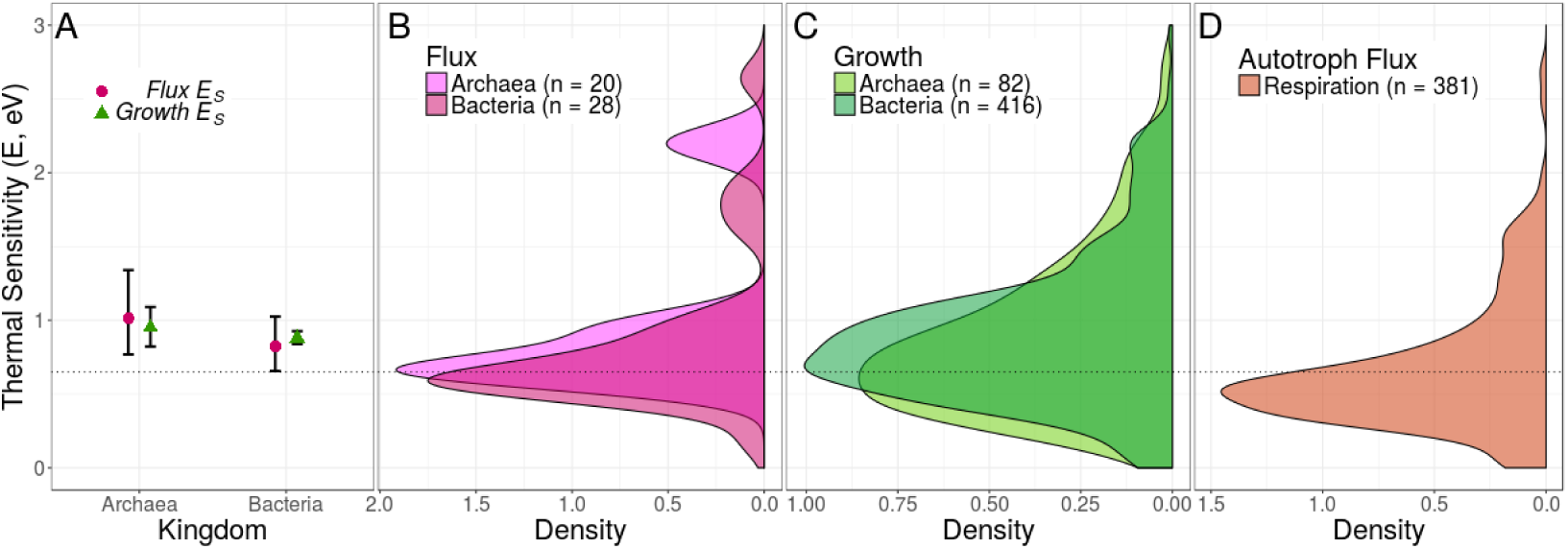
Differences in sensitivity (activation energy) of short-term thermal responses across taxonomic groups. **A.** Comparison of the intra-specific thermal sensitivity (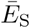) for growth and metabolic fluxes. CIs for growth rate thermal sensitivity fall within those for metabolic fluxes, and each sit above 0.65eV (dotted line) for both archaea and bacteria (bacteria growth 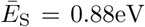, n = 416; bacteria flux 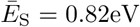, n = 28; archaea growth 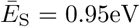, n = 82; archaea flux 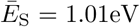, n = 20). **B.** Density plot of flux *E*_S_ values for archaea and bacteria. **C.** Density plot of growth rate *E*_S_values for archaea and bacteria. **D.** Density plot of *E*_S_ values for respiration rate TPCs in autotrophs, showing comparatively lower mean thermal sensitivity than those of the distributions for prokaryotes (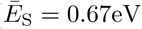, n = 381).

To further understand these findings we also categorised the data into various groups based on functional traits (Fig. 3B). Again we find mean and median thermal sensitivity *>*0.65eV in the majority of functional groups, suggesting that this high *E* is a trait generally conserved across prokaryotic organisms.

Here we have focused on the TPCs (and activation energies) of population growth rate. However, to understand the implications of the short- and long-term thermal responses of prokaryotes for ecosystem functioning it is necessary to test whether these reflect the activation energies of underlying metabolic flux rates. To investigate this, we assembled another thermal response dataset (Methods) for metabolic fluxes recorded in prokaryotes and asked whether, on average, thermal sensitivity is equivalent for growth rate and metabolic fluxes. We find that average intra-specific *E* values for growth rate TPCs were similar to, and statistically indistinguishable from the mean activation energy for metabolic fluxes (bacteria flux 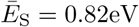; archaea flux 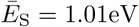; Fig. 4A, see supplementary table S1 for a list of fluxes analysed).

Furthermore, we compared both the prokaryotic growth rate and flux *E*_S_ distributions, with yet another dataset (Methods) on thermal sensitivity of respiration in autotrophic eukaryotes. The results (Fig. 4D) further support a lower thermal sensitivity of short term responses for eukaryotes than prokaryotes (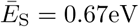 with CI = 0.63 – 0.72, median = 0.57).

### Potential Ecosystem-level Impacts

Our results higher sensitivity of both short- (higher intraspecific activation energies) and long-term (higher interspecific activation energies – a HiB constraint) thermal responses in mesophilic prokaryotes may have profound implications for responses of ecosystem fluxes to climatic warming. To illustrate this, we built a simple mathematical model to calculate the potential change in the contribution of heterotrophs to ecosystem carbon efflux (Methods). Using our new estimates of 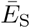 and *E*_L_ to parameterize this model, we calculate the impact of short- and long-term warming on the thermal response of carbon flux of model ecosystems that differ in composition of autotroph vs. heterotroph and eukaryote vs. prokaryote biomass. The results (Fig. 5) show that the difference in prokaryotic vs eukaryotic thermal sensitivities can substantially change the predicted increase in carbon efflux due to warming on the short- as well as long-term. For example, compared to the case where both prokaryotes and eukaryotes have the same short-term thermal sensitivity of 0.65eV (the assumption made by most current ecosystem carbon flux models ^32,33,34^), using the actual difference in sensitivity that we have found (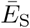 0.65eV for eukaryotes vs. 0.87 for mesophilic bacteria; Table 1; Figure 2), the flux increases by *∼*8% with 10°C short-term warming for a ecosystem composition of 50% heterotrophs (50% of which in turn are bacteria). This calculation based on the average intra-specific activation energy is relevant to short-term increases in ecosystem fluxes without evolution or acclimation in response to, for example, temperature fluctuations from time-scales of minutes to days (10°C is at the upper end of daily temperature fluctuations that organisms may typically experience ^35^). When we consider the effects of longer-term warming (such as through gradual global climate change) on the prokaryotic sub-community using the inter-specific (evolutionary) thermal sensitivity, *E*_L_ (0.98eV), we find that modelled ecosystem flux increases by *∼*5% with 4°C warming (again with 50% heterotrophs of which 50% are bacteria) compared to a baseline where the long-term thermal sensitivity is 0.65eV for all components of the ecosystem. The actual increase in flux may indeed be higher, but is dependent upon the ratio of prokaryotic biomass to eukaryotic biomass within the ecosystem, a quantity for which estimates vary widely ^8,13,36,37,38^. In our model, each percentage point increase in prokaryotic biomass within the heterotrophic component causes a flux increase of 0.05*−*0.15%, depending on the quantity of prokaryotic biomass already in the system.

**Figure 5:**
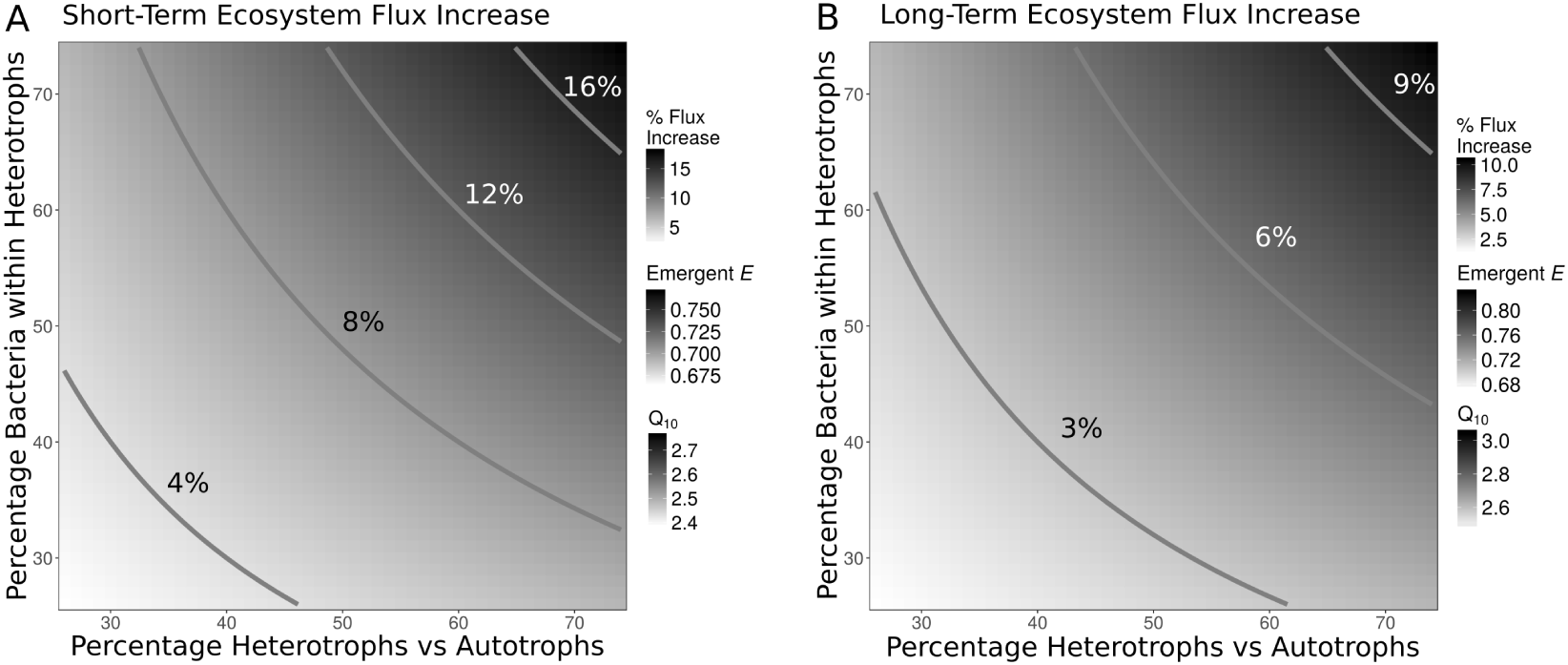
Potential changes in climate-driven short- and long-term ecosystem carbon flux due to difference in sensitivity between prokaryotic and eukaryotic thermal responses. **A.** Heat map of % short-term increase in flux with 10°C temperature increase of model ecosystems with bacteria having a different activation energy on average than eukaryotes, relative to ecosystems with all components having the same (0.65eV) average activation energy. The flux change is shown over a range of ecosystem biomass compositions in terms of heterotrophs vs. autotrophs and bacterial proportion of the heterotrophs. The scale of emergent activation energies and *Q*_10_s for the ecosystems with amplified flux are also shown. **B.** Similar to A, but for long-term flux increase under a 4°C warming scenario. Values for the short- and long-term thermal sensitivity of bacterial thermal responses used in these calculations are our estimated 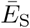 and *E*_L_ respectively for mesophilic bacteria (Table 1). The mathematical model is described in Methods.

## Discussion

Our finding of high (relative to eukaryotes) intra-specific thermal sensitivities (activation energies, *E*_S_) in prokaryotes is consistent with previous work on methanogenic archaea ^17^ and cyanobacteria ^39^, but has never been demonstrated across all major lineages of prokaryotes. In particular, Yvon-Durocher *et al.* ^17^ have argued that the high methanogen *E*_S_ are expected to translate into an increased ecosystem-level methane production at longer temporal and spatial scales. Our results suggest how these two different scales of response may be related — the short-term responses may be coupled with a Hotter-is-Better constraint which results in the flux at thermal optimum also increasing with (longer-term) adaptation. Moreover, this coupling across timescales is expected not just in methanogens, but across most major mesophilic prokaryotes, including those involved in aerobic respiration. The data do not allow us to determine the time-scale of the adaptation resulting in the HiB pattern, but numerous previous studies have shown rapid adaptation of prokaryotes to experimental warming conditions ^40,41,42^.

Due to this adaptive capacity, as global temperatures rise prokaryotes would be expected to respond to new environmental temperatures rapidly, in effect pushing them further along the global (inter-specific) HiB curve (Fig. 1A). Alternatively, species sorting may occur such that prokaryotes inherently better-adapted to higher temperatures take advantage of temperature increases. This would have the same overall effect because these prokaryotes would also effectively be further up the inter-specific temperature response curve (Fig. 2). In either case, under HiB, we can expect global warming to result in prokaryotic communities with higher metabolic rates on average. Thus overall, our results suggest that further production of greenhouse gases from the prokaryotic component of ecosystems is likely to increase in general, and at a greater rate than that by component eukaryotic organisms (Fig 5).

While in general, we see a tendency towards high thermal sensitivity (*E*_S_) in prokaryotes, there are taxonomic subgroups within our dataset for which this is not the case (Fig. 3). For example, 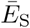 for mesophilic archaea as a whole does not deviate significantly from the MTE 0.65eV average (Table 1). This is largely because this subgroup is primarily comprised of strains from *Halobacteria*, which have thermal sensitivities significantly lower than 0.65eV (Halobacteria 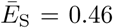; CI = 0.38 – 0.58; fig 3A). This is likely a result of their extremophilic niche imposing unusual constraints on their physiology (these archaea have been isolated only from high salinity lakes). In general, it may be harder to make generalisations about short-long-term thermal responses across taxa for archaea as a whole, because these prokaryotes are partly typified by their propensity to adapt to different types of extreme environments^43^. We also note that while the majority of heterotrophic bacteria in our dataset respire aerobically, there are a number of anaerobic strains, the majority of which were grown under various fermentation conditions. However, when we consider these groups of bacteria separately, we see no significant difference between their mean intra-specific thermal sensitivities (aerobic 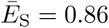, CI = 0.81 – 0.91, n = 221; fermentation 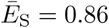, CI = 0.77 – 0.96, n = 62). Ultimately, despite all this variation, we find that both, the short-term (intra-specific) and long-term (HiB hypothesis) amplification of metabolic rate holds true for the mesophiles (≲ 45°C), a temperature range in which most of the biomass on the planet exists.

For simplicity, when parametrising our ecosystem model we used 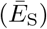 and *E*_G_ calculated from all of the mesophilic bacteria in the dataset, as we expect a huge amount of variation in the taxa present at the ecosystem scale. To expand this work further it may be possible to consider more specific situations where certain prokaryotes may dominate in certain environments based on global biogeographic studies ^44^. However, the majority of microbial taxa are known only from sequencing data^45^, for example *Acidobacteria* are thought to make up in the region of 50% of soil biodiversity ^46^, yet very few strains have actually been cultured and therefore have TPCs available ^47^. Thus in practice it may not be feasible to accurately parametrise this sort of model based on patterns of microbial biogeography and therefore, using a global average is appropriate.

We have focused on the ecosystem consequences in the face of global change, but our results also have implications for understanding of prokaryotic physiology. We are not aware of any previous work showing that prokaryotes differ systematically in their thermal sensitivity from eukaryotes. Therefore, further studies are needed to explore the mechanistic basis of this difference, and may reveal a major physiological transition mediated by an increase in cellular complexity as well as multi-cellularity in eukaryotes ^48,49^. Also, our comparisons for growth rate and metabolic flux *E* are simply averages across strains. Direct within-strain comparisons of growth rate (a slower thermal response) and the more instantaneous metabolic flux TPCs will be needed in order to fully understand the coupling of positive intra-specific and inter-specific thermal responses we have found here.

In summary, our results significantly deviate from current assumptions about the thermal sensitivity of heterotrophic respiration in ecosystems, and should be considered in ongoing efforts to model the impacts of climate change on ecosystem fluxes. More work needs to be undertaken to address whether intra-(short-term) and inter (long-term)-specific thermal responses are similarly conserved across other groups of organisms that are important for ecosystem function, such as fungi and insects in terrestrial, and phytoplankton and zooplankton in marine ecosystems.

## Methods

### Data Collection

We compiled a dataset of published prokaryotic thermal performance curves (TPCs) by searching the literature for papers with these data and using digitisation software to collect the thermal performance point estimates. Candidate TPC data was identified via manual searches of google scholar and pubmed databases. Search terms such as ‘bacteria’, ‘bacterium’, ‘archaea’, ‘archaeon’, ‘temperature’, ‘temperature response’, ‘thermal response’, ‘growth’, ‘adaptation’, were used to find papers with response data particularly for growth rates. Later searches included terms such as ‘characterisation’, ‘isolation’, ‘nov.’, ‘novel’, ‘gen.’, ‘sp.’, as it became clear that thermal responses were often tested in publications describing newly isolated species/strains. When presented as a response curve figure, ‘Plot Digitizer’ software ^50^ was used to extract data points, including error bounds when reported. The ‘Taxize’ R package ^51^ was used to standardise taxonomy of extracted data to the NCBI database. The papers were also manually searched to collect data on growth conditions as well as other metadata where possible (historical lab growth conditions, sampling location). In instances where doubling rates or doubling times were reported, we used Doubling time *t_d_* = ln(2)*/µ* to calculate the maximum specific growth rate. Raw data were normalised to rates per second and degrees Celsius for use in modelling comparisons. In total we collected 542 prokaryotic growth rate TPCs.

Although we primarily collected growth rate data as a measure of fitness in order to test HiB, we additionally collected 54 TPCs covering various metabolic fluxes for comparison to growth rate TPCs. Our complete prokaryote dataset comprises 596 TPCs from 482 unique prokaryote strains across 239 published studies.

Finally, we compiled thermal response data for respiration rates in autotrophs from the literature using the same methods for digitisation and data collation as for the prokaryote dataset. In total this autotroph dataset comprises 381 respiration rate TPCs from 140 unique autotroph species (98 vascular plants, 4 mosses, 11 green algae, 22 red algae and 5 brown algae species).

### Biological replicates and pseudoreplicates

We use prokaryotic “strains” to designate separate prokaryotic taxonomic entities with potentially differing TPCs. If a single study provided multiple TPCs from the same prokaryotic strain under the same conditions, these were considered pseudoreplicates. In these cases, all data were collected and a single Sharpe-Schoolfield fit was computed for the combined set of points, yielding a single set of TPC parameters. Where multiple TPCs were provided for the same strain under different growth conditions, these were considered as separate biological replicates, however in practice this is only the case for two replicates in each of two different strains in our dataset. Where TPCs were obtained from prokaryotes identified only to the species level (or higher), these were considered biological replicates as likely representing different strains of those species.

### Model Fitting

To each TPC in the dataset, we fitted a modified Sharpe-Schoolfield model ^52^ (eq. 1):

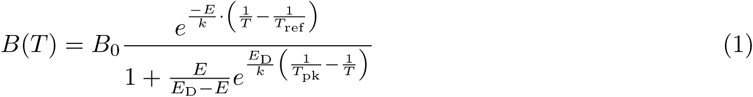

Here, *T* is temperature in Kelvin (K), *B* is a biological rate, *B*_0_ is a temperature-independent metabolic rate constant approximated at some (low) reference temperature *T*_ref_, *E* is the activation energy in electron volts (eV) (a measure of “thermal sensitivity”), *k* is the Boltzmann constant (8.617 *×* 10^−5^ eV K^−1^), *T*_pk_ is the the temperature where the rate peaks, and *E*_D_ the deactivation energy, which determines the rate of decline in the biological rate beyond *T*_pk_. We fit this model to individual TPCs and solve for *T* = *T*_pk_ to calculate the population growth rate at *T*_pk_ (*P*_pk_) for each strain. Note that this has been reformulated from the model presented in the original paper, to include *T_pk_* as an explicit parameter ^53^.

Each strain’s TPC has a potentially different *T*_pk_ and *P*_pk_. Compiling these values across strains yields an inter-specific thermal response curve (Fig 1). We fit the Boltzmann-Arrhenius equation (eq. 2, essentially the numerator in eq. 1) to these peak values to calculate inter-specific activation energy.

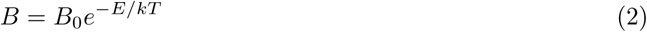

All Boltzmann-Arrhenius and Sharpe-Schoolfield model fitting was performed in Python with the NumPy package, using a least squares regression method to minimise the fits.

### Comparing Short- and Long-term thermal responses

We determined whether Hotter-is-Better by testing whether the activation energies from intra-(short-term) and inter-specific (long-term) TPCs were (statistically) significantly different. For each intra-specific curve we fitted the Sharpe-Schoolfield model (eq. 1) and extracted the intra-specific activation energy (*E*_S_), peak temperature (*T*_pk_) and corresponding growth rate (*P*_pk_) for each curve. TPCs without a peak are thus excluded from this analysis.

To estimate *E*_L_ we fitted the Boltzmann-Arrhenius model (eq. 2) to the *T*_pk_ and *P*_pk_ values estimated from the intra-specific thermal responses. To account for uncertainty in the original Sharpe-Schoolfield model fits to the intra-specific curves, we fitted Boltzmann-Arrhenius using a weighted regression (see accounting for uncertainty). In order to provide a comparison between intra- and inter-specific activation responses, we used bootstrapping to generate confidence intervals (CIs) around the mean in each case. To provide boostrapped CIs for *E*_L_ from the modified Boltzmann-Arrhenius fits, the data was re-sampled with replacement 1,000 times, with the model re-fitted to this data each time and the CIs defined as the U.S. ^th^ and 97.5^th^ percentiles of *E* values extracted from these fits. 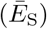 was calculated as the weighted mean *E*_S_ for the group (see supplementary methods), and CIs were taken as the 2.5^th^ and 97.5^th^ percentiles from the resultant distribution of *E*_S_ values from a bootstrap of the weighted mean.

We then determined whether the data was consistent with either of the three hypotheses (main text Fig. 1) by comparing the overlap of confidence intervals of the relevant *E* estimates. First, we tested whether 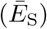 was greater than zero (null hypothesis that the CI includes zero). Second, we tested whether *E*_L_ was greater than zero (null hypothesis that the CI includes zero). Finally, if both 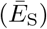 and *E*_L_ were positive, we tested whether they were significantly different to each other (null hypothesis, that the CIs for 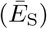 and *E*_L_ don’t overlap). Under a HiB scenario, *P*_pk_ will increase with *T*_pk_ across strains, and according to MTE this is best quantified by a Boltzmann-Arrhenius model. As a result, the Boltzmann-Arrhenius activation energies from the intra- and inter-specific responses should be positive and any differences between them not statistically significant, i.e. the confidence intervals of 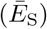 and *E*_L_ should overlap each other, but not zero. Alternatively, if growth rates are not constrained by thermodynamics and *P*_pk_ does not increase with temperature, then *E*_L_ will be close to zero (CI for *E*_L_ includes zero), and HiB can be rejected. Finally, in scenarios where thermodynamic constraints may be partially evident but somewhat overcome by adaptation, 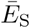 and *E*_L_ will both be positive, but with 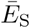 being significantly greater than *E*_L_ (i.e. 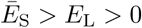)

### Accounting for Statistical Uncertainty

Weighted means were used to account for uncertainty in Sharpe-Schoolfield point estimates when calculating 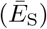 and when fitting inter-specific Boltzmann-Arrhenius curves. After performing Sharpe-Schoolfield fits we extracted the *E* and *µ*_pk_ point estimates as well as the covariance matrix. We then sampled 1,000 times from a bivariate distribution accounting for the covariance, producing 1,000 model parameter combinations. We used these parameters to generate 1,000 different Sharpe-Schoolfield curves, providing a distribution of *E* and *µ*_pk_ from which we took the standard deviations (*SD*_E_ and *SD_µ_*) as a measure of uncertainty. In some cases the Sharpe-Schoolfield fit did not produce a covariance matrix and these fits were excluded from further analysis.

When combining *E* values across strains to calculate 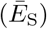 we, took weighted arithmetic means of *E*to account for uncertainty in the original fits, where *W eight* = 1*/*(*SD*_E_ + 1). Similarly, when fitting Boltzmann-Arrhenius, we apply a weighting to *µ*_pk_ where *W eight* = 1*/*(*SD_µ_* + 1).

Applying these weightings does not alter the main results we obtain from this study in terms of whether the HiB hypothesis is accepted or not for different groupings, however we felt that it was important to acknowledge and account for error in the underlying Schoolfield fits so that our results were not skewed by poor parameter estimates from questionable fits, hence this step was included. Figure S2 illustrates the differences between 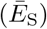 calculated with and without a weighting – applying a weighting pushes 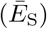 down a little, likely due to high *E* values obtained from fits to lower quality data. In either case, with or without a weighting, 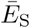 falls significantly above the 0.65eV MTE average activation energy for both Bacteria and Archaea.

### Taxonomic and Physiological Groupings

Pychrophiles and mesophiles inhabit low to medium temperature ranges, while thermophiles and hyper-thermophiles grow at much higher temperatures^54^. The distinction between mesophiles and thermophiles is usually defined relatively arbitrarily, with mesophiles often considered strains with thermal optima up to 45°C and thermophiles those with thermal optima of 55°C and above ^54^. Corkrey *et al.* ^28^ found a peak in microbial growth rates at *∼*42°C (mesophile peak) followed by an attenuation of maximum growth rates until a second peak at *∼*67°C (thermophile peak), suggesting a biological transition between mesophiles and thermophiles.

In order to determine whether it was appropriate to consider mesophiles and thermophiles separately, we performed a break-point analysis on our dataset using the ‘Segmented’ R package ^55^. Segmented is not compatible with non-linear least-squares (nls) fitting, so this was performed with a linearised version of Boltzmann-Arrhenius, i.e. *x ∼ y* where *x* = 1*/*(*kT*_pk_) and *y* = *log*(*µ*_pk_). As this process was merely to confirm whether it was appropriate to split the data into mesophiles and thermophiles as suggested by eye, it is not important that these linearised fits may give slightly different slope and intercepts to the weighted nls fits. Using this methodology we determined significant break-points for bacteria and archaea within our growth rates dataset at 40.48°C and 46.21°C respectively. These are similar to the *∼*42°C mesophile growth rate peak seen by Corkrey *et al.* ^28^ and were thus used as cut-off points for defining mesophiles and thermophiles in our analysis.

In addition, archaea are typified by their adaptations to energetically demanding niches, while in contrast bacteria perform better in more “ambient” environments^43^. A major physiological difference between these taxa lies in their fundamentally divergent membrane structures. This affects these organisms’ abilities to maintain proton gradients and thus drive metabolism under different conditions ^43^, a difference that may be particularly important for thermal performance. As such, we separate bacteria and archaea in our analysis as disparate organisms with divergent evolutionary histories.

In order to classify prokaryotes by the energy generating metabolic processes that they use, we took note of the growth conditions used when initially digitizing the TPC data. For the majority of heterotrophic bacteria and archaea this was simply whether they were grown under aerobic or anaerobic (fermentative) conditions. However there are also a number of strains utilising more exotic metabolic processes such as methanogenesis, sulfur reduction, etc. In these cases we matched taxa against those able to utilise certain metabolic reactions according to Amend & Shock ^56^ before manually checking the culture conditions in each study for the metabolites required for certain metabolic processes.

We also categorised taxa by their status as potential pathogens. We matched taxon names against the database of host-pathogen interactions provided in Wardeh *et al.* ^57^ to understand whether each strain was potentially pathogenic, and what taxa they were known to infect.

### Ecosystem carbon flux model

To quantify the effect of differences in activation energy of respiration between prokaryotes and eukaryotes on carbon flux, one can calculate the percentage increase in flux (*F_x_*) of an ecosystem as,

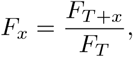

where *x* is the temperature increase (at the end of a warming scenario), and *F_T_* and *F_T_* _+*x*_) are the fluxes at the two temperatures. Because ecosystem carbon flux at night (i.e. without photosynthesis) is the sum of autotrophic and heterotrophic respiration rates weighted by the biomasses of these compartments, we can re-write *F_x_* as:

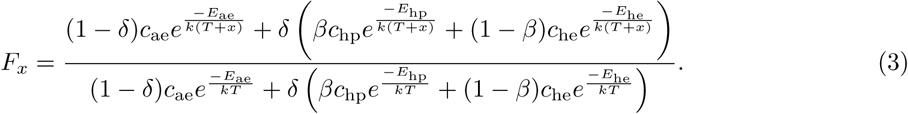

Here, each compartment’s total flux contribution (identified by a subscript: autotrophic eukaryotes = ae, heterotrophic prokaryotes = hp; heterotrophic eukaryotes = he) is modelled as a Boltzmann-Arrhenius equation, with *c* a normalisation constant. Each compartment’s contribution is weighted by the biomass proportionality constants: *δ* is the proportion of heterotrophic biomass in the ecosystem, while *β* is the proportion of prokaryotic biomass within the heterotrophic component (so 1 *− β* is the proportion of non-prokaryotic heterotrophs such as fungi or insects). We do not use the Sharpe-Schoolfield model here because it does not apply to long-term thermal responses (Fig. 1), whilst for short-term responses most warming as well as temperature fluctuations are expected to occur within an “operational temperature range”, which excludes temperatures greater than *T*_pk_ (the heat-stress region) ^58^. We do include any potential contribution of autotrophic prokaryotes (such as cyanobacteria), as these are not expected to provide a significant flux contribution to a typical terrestrial ecosystem.

We then use eqn. 3 to calculate the percent change in ecosystem flux due to differences in activation energies of the three compartments (*E*_ae_, *E*_hp_, and *E*_he_):

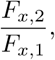

where *F_x,_*_2_ and *F_x,_*_1_ are the warming-induced flux changes in ecosystems with and without differences in activation energies of the compartments, respectively (the value of the heat map in Fig. 4). That is, for *F_x,_*_1_, all *E* values, i.e. *E*_au_, *E*_hp_ and *E*_he_ in Eq. 3 = 0.65eV. This is the assumption made by most current ecosystem carbon flux models ^32,33,34^. For *F_x,_*_2_, the differing activation energies were parametrised using either the mean of the estimated *E*s for the short-term (intra-specific) or the long-term (inter-specific) TPCs (Table 1; Fig 2). For this we used estimates of *E*_L_ and 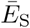 from mesophilic bacteria (long-term “evolutionary” *E*_L_ = 0.98eV, short-term “instantaneous” 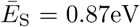) only, because the archaea in our data are largely composed of strains adapted to extremophilic niches, which are largely irrelevant from a global warming perspective.

We calculated the emergent *E* of the *F_x,_*_2_ ecosystems (flux response to warming when prokaryotic and eukaryotic thermal sensitivities differ), which is the the average of activation energies for each ecosystem compartment weighted by its biomass proportion:

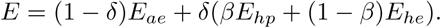

We also calculated the emergent *Q*_10_ of the *F_x,_*_2_ ecosystems, as it is a widely used measure in climate change models of carbon flux ^33,59^:

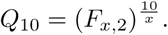

We chose a warming magnitude *x* = 10°C for short-term responses because this at the upper end (e.g., generally, at higher latitudes) of the range of daily (over 24 hrs) fluctuations that organisms experience ^35^. For long-term warming scenarios, we used *x* = 4°C, the approximate upper end of the range for the year 2100 projected by the IPCC^60^.

The biomass proportions *δ* and *β* were varied to capture the effect of different ecosystem compositions. In a typical forest ecosystem, the contribution of autotrophic to heterotrophic (mostly soil) respiration has been estimated to be approximately 50% each ^36^. This heterotrophic component would be comprised largely of prokaryotes and soil fungi biomass, the ratios of which have shown to vary widely depending on soil type and the experimental methodology used ^13^. Here we vary the percentage of heterotrophs within an ecosystem (*δ*) between 25 – 75% and the percentage of prokaryotes within heterotrophs (*β*) between 25 – 75% to generate a range of potential scenarios in Fig. 5.

## Author Contributions

TPS and SP conceived the study. TPS, TJHT, SS and GY-D compiled the data. TJHT wrote the thermal model fitting code. TPS and TJHT wrote the analysis code. TPS, TJHT and SP analysed the data. TPS and SP wrote the manuscript, and all authors contributed to the revisions.

## Acknowledgements

We would like to thank Rebecca Kordas for helpful comments on an earlier version of the manuscript. TPS was supported by a BBSRC DTP scholarship (BB/J014575/1). BGC, SS, and SP were supported by a NERC grant awarded to SP (NE/M004740/1). TB was supported by an ERC starting grant (311399-Redundancy). GY-D was supported by an ERC starting grant (677278 TEMPDEP).

