## Supplementary Information for "Metabolic rates of prokaryotic microbes may inevitably rise with global warming"

#### Contents

**1 The Association Between Thermal Optimum and Culturing Temperature 1**

**2 Comparison of Intraspecific Thermal Sensitivities 2**

**3 Microbial Metabolic Fluxes 3**

#### 1 The Association Between Thermal Optimum and Culturing Temperature

Here we show that there is strong adapatation of prokaryotic growth rates to their evolutionary thermal regime. From the information collected for each thermal response curve, we extracted data on the routine culturing conditions of each strain prior to the experiment. It is often stated that organisms tend to operate at temperatures below their peak temperature allowing a buffer before temperature becomes detrimental to fitness, and this is borne out in our data with peak growth temperature ($T_{\text{pk}}$) tending to be marginally higher than the standard laboratory growth temperature ($T_{\text{lab}}$) (Fig. S1). When organisms were provided by a cell culture bank, this temperature is simply the temperature that the strains were cultured at by the experimentalists prior to use, however in studies of novel organisms, this temperature was often also the temperature that was used during isolation of the organism. This may hint at rapid adaptation by these organisms to the routine culture conditions, or in the case of isolation experiments, possibly species sorting such that species isolated were simply those with the appropriate thermal tolerance range. In the high temperature range we see that in Bacterial strains $T_{\text{pk}}$ begins to tend to fall below $T_{\text{lab}}$, potentially indicating that this temperature range is past the limit of thermal adaptation for Bacteria - i.e. despite being routinely cultured at high temperatures, these strains are unable to adapt to these temperatures due to the inherent biological properties of Bacterial cells.

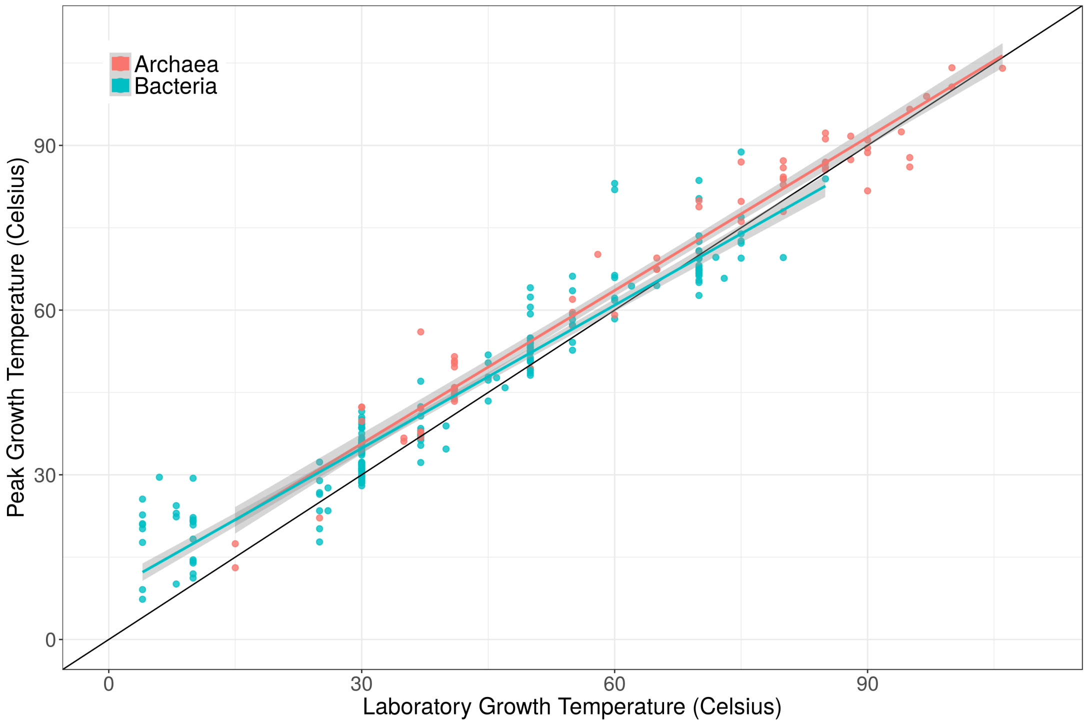


Figure S1: Peak growth temperature ($T_{\mathrm{pk}}$) against the laboratory growth temperature ($T_{\mathrm{lab}}$), with linear models fit for bacteria and archaea (bacteria intercept = 8.79, slope = $0.87 \pm 0.04$, $R^2 = 0.91$, $p < 0.00001$; archaea intercept = 7.77, slope = $0.93 \pm 0.05$, $R^2 = 0.96$, $p < 0.00001$) and a 1:1 line shown (black). In general there is a strong association of $T_{\mathrm{pk}}$ with $T_{\mathrm{lab}}$. The confidence intervals for the slopes are below 1 however, indicating that prokaryotes tend to be unable to adapt to very high culturing temperatures.

### 2 Comparison of Intraspecific Thermal Sensitivities

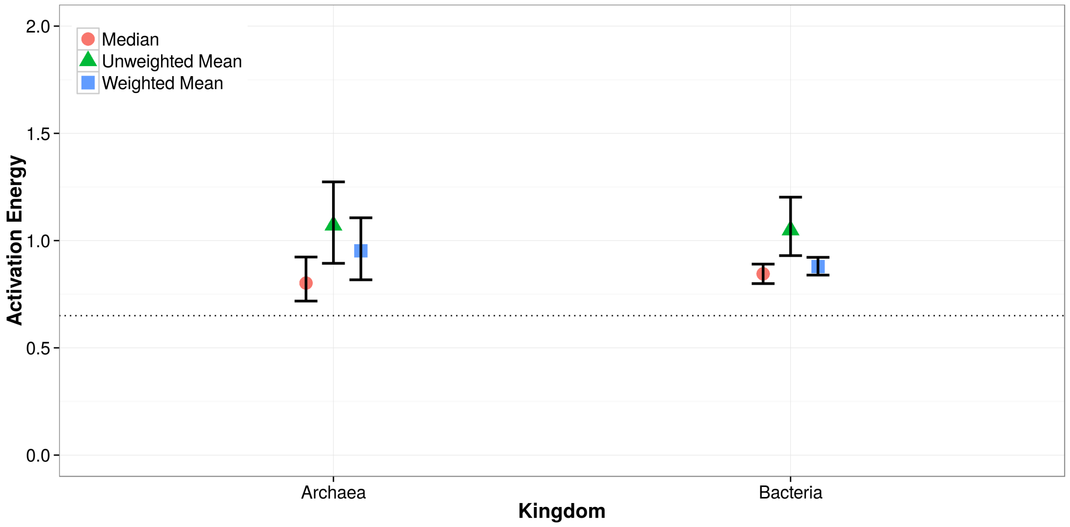


Figure S2: Comparison of intra-specific thermal sensitivities ($E_{\mathrm{S}}$) calculated as the median, unweighted and weighted means respectively. The median is lower than the mean $E_{\mathrm{S}}$ for both Bacteria and Archaea as expected due to the right-skew of the data[1], however in all cases $E_{\mathrm{S}}$ is above the 0.65eV average (dotted line) expected from the Metabolic Theory of Ecology, which we find only holds for Eukaryotes (Main text Fig. 4).

#### 3 Microbial Metabolic Fluxes

To test whether thermal sensitivity for growth rate (fitness) reflects the thermal sensitivity of underlying metabolic fluxes, we assembled prokaryotic flux data for comparison. Metabolic flux data is sparse in the literature in comparison to growth rate data, however we were able to find some temperature-dependent flux data from a number of different studies. The various metabolic flux data that we compiled for this analysis and their respective sources are detailed in Table S1.

Table S1: Metabolic fluxes which we fitted TPCs to for comparison to growth rate TPCs.

| **Metabolic Flux** | **References** |
|---|---|
| Sulfur oxidation rate | [2] |
| Fe3+ oxidation rate | [2] |
| Fe2+ oxidation rate | [2, 3] |
| Sulfide production rate | [4] |
| Methanogenesis | [5, 6, 7, 8, 9, 10, 11] |
| Nitrate production rate | [12] |
| Nitrate removal rate | [13] |
| Fe2+ production rate | [14] |
| Dehalogenation rate | [15] |
| Sulfate reduction rate | [16, 17, 18] |
| Caffeine degradation rate | [19] |
| Hydrogen sulfide production rate | [20] |
